# Mesenchymal-like glioma cells are enriched in the gelatin methacrylate hydrogels

**DOI:** 10.1101/2021.03.24.436751

**Authors:** Nameeta Shah, Pavan M. Hallur, Raksha A. Ganesh, Pranali Sonpatki, Divya Naik, Komal Prasad Chandrachari, Ralph B. Puchalski, Aditya Chaubey

## Abstract

Glioblastoma is the most lethal primary malignant brain tumor in adults. Simplified two-dimensional (2D) cell culture and neurospheres *in vitro* models fail to recapitulate the complexity of the tumor microenvironment, limiting its ability to predict therapeutic response. Three-dimensional (3D) scaffold-based models have emerged as a promising alternative for addressing these concerns. One such 3D system is gelatin methacrylate (GelMA) hydrogels, which can be used for modeling the glioblastoma microenvironment. We characterized the phenotype of patient-derived glioma cells cultured in GelMA hydrogels (3D-GMH) for their tumorigenic properties using invasion and chemoresponse assays. In addition, we used integrated single-cell and spatial transcriptome analysis to compare cells cultured in 3D-GMH to cells *in vivo*. Finally, we assessed tumor-immune cell interactions with a macrophage infiltration assay and a cytokine array. We show that cells cultured in 3D-GMH develop a mesenchymal-like cellular phenotype found in perivascular and hypoxic regions present in the core of the tumor, and recruit macrophages by secreting cytokines in contrast to the cells grown as neurospheres that match the phenotype of cells of the infiltrative edge of the tumor.

## 1. Introduction

Glioblastoma, the most common and aggressive form of primary brain tumors in adults, is characterized by angiogenesis and diffuse infiltration(1). Despite a rigorous treatment regimen, which entails surgery followed by radiation and chemotherapy, the prognosis is dismal(2). Although large-scale molecular data generated over the last two decades has provided insight into the cellular and molecular mechanisms underlying GBM pathogenesis and progression, effective treatment options have yet to be established(3). The lack of therapeutic progress can be attributed to the complex inter-and intra-tumor heterogeneity of GBM(4).

Tumor heterogeneity is widely influenced by genetic and epigenetic changes in tumor cells and by the complex network of tumor micro-environmental factors supported in the crosstalk(5). Initial characterization of the heterogeneity of GBM with bulk tissue transcriptome profiling suggests three molecular subtypes: proneural (PN), classical (CL), and mesenchymal (MES)(6, 7). Recent work on glioblastoma reported that molecular signatures of anatomic features delineated histologically (tumor periphery, infiltrating tumor, cellular tumor, hypoxic tumor, tumor vasculature) were highly conserved across patients and reflected the biological processes, pathways, cell-types, and microenvironment relevant to each feature(1). Subsequently, single-cell transcriptome studies of isocitrate dehydrogenase wildtype (IDHwt) GBMs have identified four patient-independent cell states: OPC (oligodendroglial lineage), NPC (neural progenitor lineage), MES-like (mesenchymal), and AC (astrocytic lineage)(8, 9). Wang et al. also show that IDHwt GBM cells lie on a single axis of gene signature ranging from proneural to mesenchymal phenotype. At both ends of the spectrum are glioma stem-like cells (GSCs) with proneural cells being highly proliferative (pGSCs) and mesenchymal stem-like cells (mGSCs) being quiescent with cytokine secretory phenotype(9). This conservation of molecular and cellular phenotype for a specific anatomic feature across patients makes simultaneous targeting of these features an attractive therapeutic approach and underscores the need for biomimetic *in vitro* models that most closely represent distinct *in vivo* microenvironments(1,8,9).

Glioblastoma biology is typically studied using 2D *in vitro* or *in vivo* models. Another model frequently used to study glioma stem-like cells employs serum-free conditions, which allows retention of original tumor stem cell properties compared to serum-supplemented conditions(10). In a recent study, to retain the cellular heterogeneity *in vitro*, the authors used serum-supplemented cultures in parallel with serum-free cultures as both cultures enrich specific glioblastoma cell-types -proneural and mesenchymal, respectively(11). Behnan and co-workers introduced the concept of mixed cell-culture grown in serum and growth factors, where both proneural cells and mesenchymal cells were preserved and contributed to highly infiltrative, hypoxic, and angiogenic tumors when implanted in mice(12). Although these strategies enrich various subtypes of GBM successfully, they do not imitate an *in-vivo* tumor environment. Given the spatial, molecular, and temporal complexity of the native GBM tumor, tissue engineering approaches towards the fabrication of complex *in vitro* models that most closely represents the *in-vivo* need to be established.

Scaffold-based 3D culture systems are physiologically relevant *in vitro* models as they retain the cell-cell and cell-matrix interactions similar to what is observed *in vivo*(13, 14). Various three-dimensional hydrogel models have been fabricated using both bio-mimetic and synthetic polymers. Hydrogels are attractive candidates to study glioma biology due to their high tissue-mimicking water content and tunable biochemical and physical properties. Gelatin methacrylate (GelMA) hydrogels are especially relevant as gelatin is a natural polymer derived from denatured type I collagen, the most abundant ECM protein. Importantly, specific ECM proteins can be incorporated into these hydrogels and their mechanical properties can be modulated to create a “tissue-biomimetic” cell culture system. Extensive chracterization for tunable mechanical property, porosity and diffusive properties have established 3D-GMH as an inexpensive, cell-responsive platform for modeling key characteristics associated with cancer metastasis, *in vitro*(15, 16). Further, cellular response to the stiffness of the hydrogels and biochemical cues have been shown to have an influence on glioblastoma invasion(17–19). Co-culture of glioma cells with various cell-types in GelMA hydrogels show varied tumorigenic properties(18,20,21). Given these advantageous properties of 3D-GMH, in this study, we used a 3D-GMH cell culture system with serum. Using patient-derived glioma cells, we demonstrate that cells cultured in 3D-GMH show higher invasive potential and increased chemoresistance to temozolomide (TMZ) compared to cells cultured in 2D. Transcriptome analysis revealed that our 3D-GMH system enriches for the mesenchymal phenotype and resembles the cells found in the perivascular and perinecrotic zones of glioblastoma tumors. Furthermore, gene expression signatures derived from the transcriptome of glioma cells grown in 3D-GMH show poor survival association in two large glioma atlases. Finally, we show that cells grown in 3D-GMH secrete increased levels of key cytokines and display a higher potential to recruit macrophages. In conclusion, we propose the 3D-GMH scaffold as a complementary *in vitro* system to the neurosphere cultures to study the biology of hypoxia-dependent and independent mesenchymal glioma cells, including understanding tumor-immune cell interactions.

## 2. Materials and Methods

### 2.1 Ethics statement

Glioma samples were collected at the time of surgery with informed consent from patients at Mazumdar Shaw Medical Center (MSMC). All the procedures were performed in accordance with recommended guidelines and approval of the Institutional Ethics Committee, NHH/MEC-CL-2014/219, Bengaluru, India.

### 2.2 Glioma patient-derived cultures

Surgically resected glioma samples were collected and transported to the laboratory on ice within one hour of the surgery. The samples were washed thrice with ice-cold phosphate buffered saline (PBS, pH 7.4) (10010023; Gibco, USA) with 1% Penicillin-Streptomycin antibiotics (P4333-Sigma-Aldrich, USA) and minced into small pieces with a scalpel before being digested with Accutase (A6964-Sigma-Aldrich, USA) for 20 minutes. The digested tissue was dissociated by pipetting up and down before passing through a 70μm cell strainer. Strained cells were washed with ice-cold Dulbecco’s Modified Eagle Medium F12 media (DMEM F12; 11320033-Invitrogen) and centrifuged twice at 1100rpm for 10 minutes at 4°C. The cell pellet was suspended in DMEM F12 media with 10% fetal bovine serum (FBS) (RM10409; HiMedia, USA) and cultured in 2D. Cell culture media was changed every 2–3 days, and cells were passaged when they reached 70-80% confluence. Cells were passaged in 2D 1-4 times before they were used for various experiments.

### 2.3 Cell line

Human glioblastoma cell line U-251MG was cultured in Dulbecco’s Modified Eagle Medium (DMEM-HG) (11995-065; Gibco, USA) and supplemented with 10% FBS (RM10409; HiMedia, USA) and with 1% Penicillin-Streptomycin (15140122-Gibco) at 37°C with 5% CO2. U937 cells (a monocyte-like cell line) were cultured in RPMI-1640 media (AL060A; HiMedia, USA) containing 20% FBS (10270106-Gibco, USA) and 1% Penicillin-Streptomycin (15140122-Gibco).

### 2.4 Neurosphere

Both cell line and patient-derived cells were initially passaged in 2D before they were propagated as neurospheres (GSC) in ultra-low attachment plates with serum-free Neurobasal™ Medium (NBM; 21103049; Gibco, USA) with 10 ng/ml of EGF (PHG0311; Gibco, USA) and bFGF (13256029, Gibco, USA), supplemented with N2 (17502048, Gibco, USA), B27 (17504044; Gibco, USA), and L-Glutamine (25030-081; Gibco, Brazil) as previously described(22). All the reagents were supplied by Invitrogen, USA, unless otherwise stated.

### 2.5 Immunofluorescence Assay

Patient-derived cells were fixed with 4% PFA for 15 minutes, blocked with 3% BSA in PBS with 0.1% Tween20 for 1 hour and immunostained with rabbit anti-nestin (1:200, PA-1-86334; Invitrogen, USA), rabbit SOX2 (1:200, PA-1-16968; Invitrogen, USA), and mouse anti-GFAP (1:200, NB300-142-A1; Novus Biologicals, USA) overnight. Following incubation, Alexa 488-conjugated goat anti-mouse (ab150077; Abcam, USA) and Alexa-594 anti-rabbit (ab150116; Abcam, USA) were added as secondary reagents. Nuclei were counterstained with Fluoroshield Mounting Medium with DAPI (ab104139; Abcam, USA). Samples were subjected to evaluation under a fluorescence microscope.

### 2.6 Fabrication of GelMA Hydrogels

GelMA hydrogels were prepared as previously described(15). Briefly, 10% (w/v) Gelatin (901771; MPBiomedicals, USA) was dissolved in PBS at 50°C for 1 hour under constant stirring, to which 20% (w/v) of methacrylic anhydride (276685; SigmaAldrich, USA) was added dropwise. Excess of PBS was added to stop the reaction and dialyzed against dH20 at 37°C, followed by freeze-drying (Alpha1-2/LDplus, Martin Christ, Germany) and storage at -20°C. Freeze-dried GelMA (10% w/v) was dissolved in PBS (10010023; Gibco, USA) along with a photo-initiator 2-hydroxy-4′-(2-hydroxyethoxy)-2-methylpropiophenone (Irgacure2959-410896, Sigma-Aldrich, USA), at a concentration of 0.5 mg/mL followed by exposure to UV-365 for 10 minutes, to obtain hydrogels. These hydrogels were equilibrated with fresh media containing 10% FBS for 24hrs prior to seeding of cells for experiments. Freshly dissociated cells were counted and seeded at a concentration of 5*10^4^ cells per well of a 24-well plate.

### 2.7 Invasion assay

Cells cultured in 2D were harvested by trypsinization, and those cultured in 3D-GMH were harvested by enzymatic degradation of the 3D-GMH using collagenase, and suspended in a serum-free basal medium. For invasion assays, these cells were then plated (10^4^ cells/chamber) on to invasion chambers (Sigma-Aldrich, 8.0μm pore size, TCP150-HiMedia) coated with 1 mg/mL matrigel (E1270-Sigma-Aldrich, USA). The upper chamber contained a serum-free basal medium. In the lower chamber, a medium with 10% FBS was used as a chemoattractant. After 24 hours, the medium was removed and the chambers were washed twice with PBS; non-invading cells were removed from the upper surface of the membrane by gentle wiping with a cotton-tipped swab; invading cells in the lower surface of the membrane were fixed with 4% formaldehyde in PBS for 10 minutes, washed twice with PBS, permeabilized with methanol for 20 minutes, washed twice with PBS, stained with 0.4% crystal violet (V5265-Sigma-Aldrich, USA) for 15 minutes, and washed twice with PBS. In each chamber, cells invaded were photographed at a magnification of 10×, and cells were counted in each field. The fold increase in invasion was calculated by normalizing the total number of invaded cells from the 3D-GMH group to the total number of invaded cells from the 2D group.

### 2.8 Chemoresponse assay

The metabolic activity of cells cultured under different conditions was analyzed by the MTT colorimetric assay. U251 cells and other patient-derived glioma cells were seeded in the 3D-GMH and 2D dishes at a density of 1 × 10^4^ per well of a 96-well plate. Temozolomide drug treatment was started after 72 hours for the cells in 3D-GMH and after 24 hours for the cells cultured in 2D. For hypoxic conditions, the respective cells in 2D were pre-incubated in hypoxic conditions, i.e., 1% oxygen for 24 hours before the drug treatment. Temozolomide (T2577-SigmaAldrich, USA) concentrations ranging from 3.9µM to 500μM were used for assaying drug sensitivity. At predetermined times, i.e., after 72 hours of drug treatment, 20μl of a 5mg/mL 3-(4, 5-dimethylthiazol-2-yl)-2,5-diphenyltetrazolium bromide (M2128-MTT; Sigma-Aldrich, USA) solution was added to each of the wells. After 2 hours of incubation, the supernatant was carefully removed and 100 μL of a dimethyl sulfoxide (ICN19605590-MP Biomedicals, USA) solution was added to dissolve the formazan crystals. After shaking for 10 minutes on a plate mini-shaker, 100μl of solution from each well was transferred into the wells of a 96-well plate, and the absorbance was read at 570nm with a reference wavelength of 690nm (Infinite F200 Pro, Tecan, USA). Four experimental replicates were averaged for each concentration per experiment. IC50 values were derived by using GraphPad Prism 5. The results of U251 and MN238 represent 3 biological repeats whereas MN298 and MN474 represent only one biological experiment, which includes 4 experimental replicates.

### 2.9 Cell-cycle analysis

U251 cells were seeded in the 3D-GMH and 2D dishes at a density of 5 × 10^4^ per well of a 24-well plate. Temozolomide drug treatment was started after 72 hours for the cells in 3D-GMH and after 24 hours for the cells cultured in 2D. And for hypoxic conditions, the respective cells in 2D were pre-incubated in hypoxic conditions, i.e., 1% oxygen for 24 hours before the drug treatment. U251 cells were retrieved from the 3D-GMH and 2D dishes after 72 hours of treatment with 3× IC50 concentrations of TMZ determined for 2D cultures in the MTT assay. The cells were fixed in 70% (v/v) cold ethanol and stored at −20°C for 1 hour, treated with RNase (10μg/ mL), and stained with 40μg/ml propidium iodide (PI; Sigma Aldrich, USA) for 30 minutes in the dark. The fractions of cells in G0/G1, S, and G2/M phase were determined by flow-cytometric analysis in a BD FACS Canto II instrument using the red fluorescence range of excited PI-stained nuclei as a measure of the DNA content. Linear displays of fluorescence emissions were used to compare cell-cycle phases and quantitate the cells.

### 2.10 RNA Extraction

Cells grown in GelMA hydrogels were dissociated using collagenase Type II enzyme (17101015, Gibco, USA) at a concentration of 1mg/ml, incubated for 2 hours at 37°C incubator, and cells grown in 2D were trypsinized to collect cell pellets. 1ml of TRIzoL (15596018, Ambion) is added to the cell pellet after a PBS wash and stored at -20°C. Frozen cell pellet was thawed on ice, and 200ul of chloroform (650498, Sigma-Aldrich, USA) was added and vortexed, and incubated at room temperature for 5 minutes. To separate the RNA into the aqueous phase, samples were centrifuged at 12000g for 15 minutes at 4°C. The aqueous phase was carefully transferred to another tube and precipitated with 500ul of isopropanol (19516, Sigma-Aldrich, USA) at RT for 15minutes. Samples were then centrifuged at 12000g for 15 minutes at 4°C. The RNA pellets were then washed thrice with 1ml of 70% ice-cold ethanol, followed by centrifugation at 7500g for 5 minutes in 4°C. The ethanol was carefully removed, and RNA pellets were dried at RT before suspending them in 40-100ul of RNAase free DEPC treated water (112420304, MP Biomedicals, USA).

### 2.11 Transcriptome analysis

MSMF: We assessed the quality of RNA with a Bioanalyzer to ensure that all samples had an RNA integrity number (RIN) of 7 or more. The construction of the RNA library was performed according to the manufacturer’s protocol using the TruSeq RNA Library Preparation Kit (TruSeq RNA Sample Prep v2, Illumina, San Diego, CA). All libraries were sequenced using Illumina HiSeq 2000 platform with 151 bp paired-end strategy. The sequenced transcriptome was aligned to the hg19 reference genome using STAR(23) algorithm with default parameters. Gene expression was quantified using the ENSEMBL reference with bedtools(24). For count data normalization, the Count per Million (CPM) method was applied using the following formula:

CPM = (count/sum (count))*1000000)

The CPM data was log2 transformed after adding a pseudocount of 1 for further analysis.

**Public domain:**

The following datasets were downloaded from the GEO database and processed for downstream analysis.

1. Patel et al. used single-cell RNA-seq to profile data from five primary glioblastomas. They further established GSC and DGC cultures for three out of the five primary glioblastomas (MGH26, MGH28 and, MGH31) to identify tumor cells with stem-like or differentiated phenotypes. The raw data were downloaded from the SRA database, which was provided in the original publication. The data processing was done using the same approach explained above.
2. Neftel et al. Generated and analyzed single-cell RNA-seq data from 28 pediatric and adult glioblastoma tumors to identify four major neoplastic cell types defined by six gene modules; 1. Mesenchymal -Hypoxia independent (MES1like) and hypoxia dependent (MES2like) mesenchymal related gene sets, 2. Astrocytic -astrocytic (AClike) marker gene set, 3. Oligo - oligodendroglial (OPClike) lineage marker gene set, and 4. Neural - stem and progenitor cell signatures (NPC1like and NPC2like) as well as two cell cycling modules namely G1S and G2M (CC). The processed data was downloaded from the broad institute single cell in Transcripts per Million (TPM) format. Recursive Consensus Clustering(25) was used to cluster the data, which resulted in a total of 80 clusters out of which 61 clusters represented the malignant cells population. For further analysis, only the malignant cell clusters were used. Neftel et al. assigned a meta-module score to each cell based on their transcriptomic profiles. Similarly, each RCC cluster was assigned a meta-module based on the frequency of modules in that cluster. The cluster was annotated with a specific meta-module if >50% of cells in that cluster belonged to the same meta-module. Each cluster was averaged across all the cells to obtain one value per gene per cluster.
3. Darmanis et al. have generated two single-cell RNA-seq datasets: one in 2015 using human adult cortical samples and another one in 2017 using four GBM patient samples. Both the datasets were downloaded and processed as mentioned in the approach above. The dataset contained a total of 4055 cells and 15 known cell types. Recursive Consensus Clustering was used to cluster the data into novel subtypes. The Darmanis data was divided into 43 clusters with 18 out of them being the subgroups of malignant cells. Single Sample Gene Set Enrichment Analysis (SSGSEA) was performed on all the cells using the Neftel meta-modules as gene sets. A cell was assigned to a specific meta-module based on its SSGSEA score. Similar cluster averaging and assignment was performed on the Darmanis clusters as mentioned above. The Neftel and Darmanis gene × cluster matrices were combined for further analysis. Only the protein coding genes were used for clustering of the single-cell datasets.
4. The Ivy Glioblastoma Atlas Project (Ivy GAP) used laser capture microdissection technique to generate molecular signatures of cells present in five major anatomic features of GBM, visible by H&E staining, i.e. leading edge (LE), infiltrating tumor (IT), microvascular proliferation (MVP), necrosis (NE), and pseudopalisading cells around necrosis (PAN). In addition to this, the atlas also contains transcriptome profiles of cells expressing community curated GBM stem cell-associated genes (ccGSC markers). These samples were grouped in three categories; CT area with the expression of one or more ccGSC markers (CT_ ccGSC), Perinecrotic zone with the expression of one or more ccGSC markers (PNZ_ ccGSC), and area around hyperplastic blood vessels with the expression of one or more ccGSC markers (HBV_ ccGSC). The ccGSC markers and the number of samples in each ccGSC histology is shown in the table 1. The CT_control samples are the CT region with the absence of one or more ccGSC markers. The expression values were averaged across all the histological features to get one value per feature per gene.
5. Evrard et. al. generated GSC and DGC microarray data from three GBM patients (SRB1, SRA5, and SRC3). The DGC cell line was established from SRC3 GSCs. CEL files were downloaded and further processed using the limma and affy packages in R. Robust Multiarray Average (RMA) was performed to obtain log2 normalized intensities of each probes. Each of the probe names were matched to associate with RefSeq accession based on the annotation provided by GPL16686 platform. Refseq accession was further converted to gene symbol. Gene level expression matrix was then created by mapping probe names to gene symbols and taking median expression values of genes if mapping to multiple probes. Paired t-test and fold change calculations were performed to obtain significant genes between GSC and DGC.
6. Ishida et. al. generated GSC and DGC microarray data for two (NCH644 and NCH690) GBM samples. The processed data was downloaded from the GEO database and the probe names were matched to associate with the gene symbols. Further differential analysis was done using the single sample comparison method.
7. TCGA and CCGA count data were downloaded from their respective data portals. The count data were log2 CPM normalized as mentioned in the methods above.

**Table 1:**
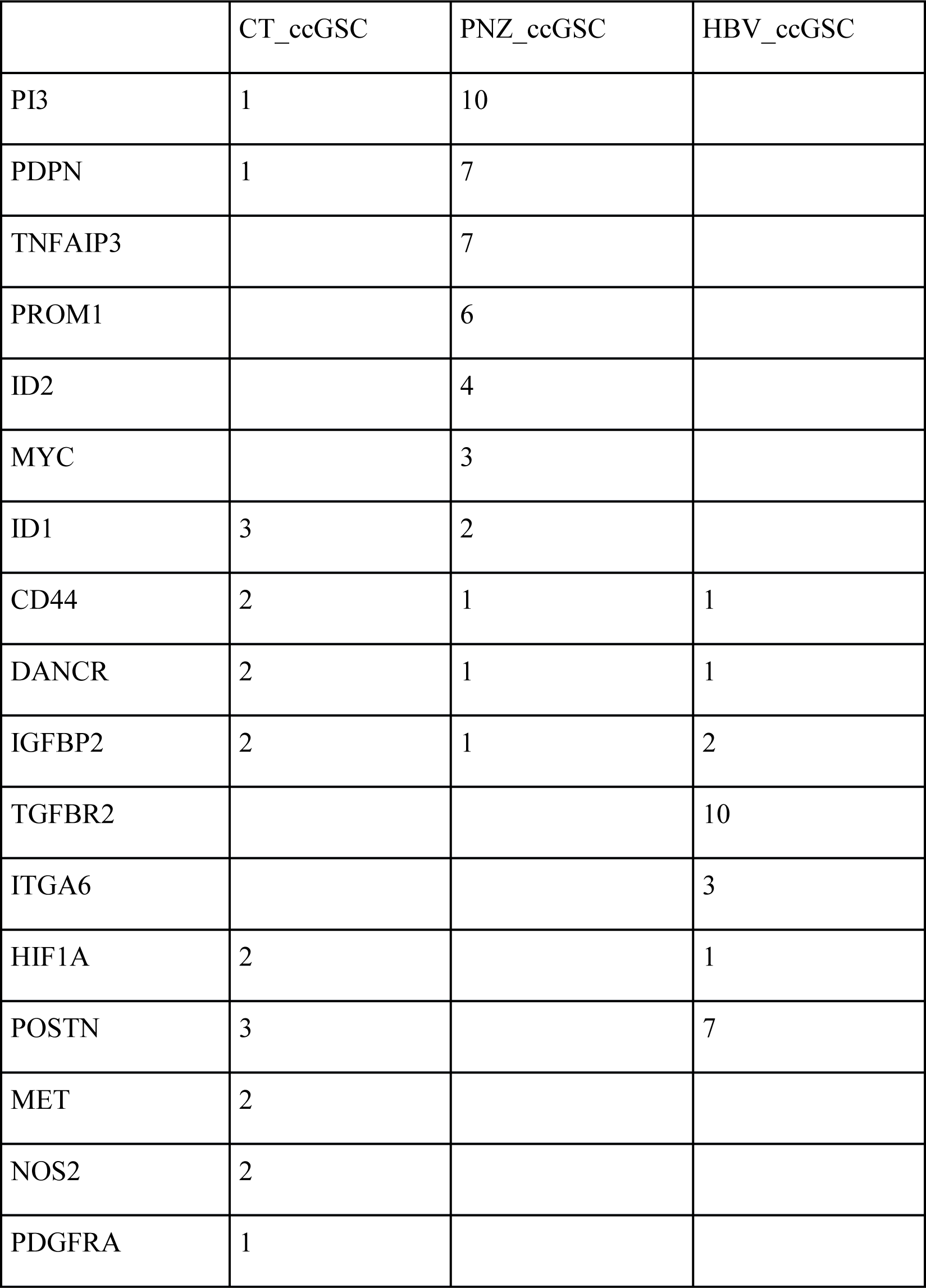
ccGSC markers and the number of samples in each histological feature

**Differential gene expression analysis: Single sample comparison:**

Differential expression analysis was performed for the following four comparisons: GSC vs. DGC, 3D-GMH vs. 2D, 3D-GMH vs. hypoxia, and hypoxia vs. 2D. As the replicates for each sample were not generated the following method was used for identification of differentially expressed genes. The linear regression model was applied to the log2 normalized gene expression values. The 99% prediction interval was then calculated for the linear regression model using the range of values from the control sample. Similar to the confidence interval, the prediction interval gives uncertainty around a single value point. For each gene, based on the linear regression model, the value of the gene in the test sample was predicted based on the value of the gene in the control sample at a 99% prediction interval. We used the *predict()* function in R to do so. A gene was considered to be upregulated/downregulated if:

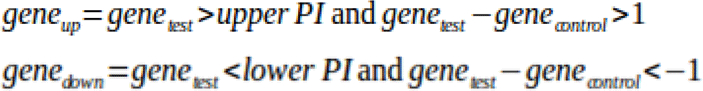

The z-score for each gene was calculated using the following formula:

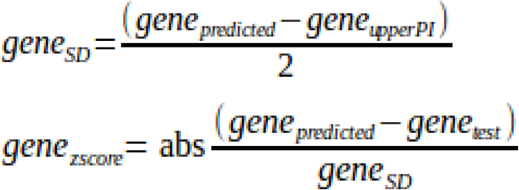

The differential gene expression data was plotted using the volcano plots (Supplementary Fig.-S3) in R. For the volcano pots, z-score was used as y-axis instead of -log10 of p-value in case of single sample comparisons.

Using the single-sample comparison method, differential gene expression was calculated across four groups. The 3D-GMH list was constructed by taking genes that showed overexpression in all the three comparisons (3D-GMH vs. 2D, MN161, MN285, and MN474) and the overexpression was statistically significant in at least two of the comparisons. For the GSC list, we used data from four different studies(10,26–28). A gene was part of the list if it was overexpressed in two comparisons in the Patel et. al. dataset or was overexpressed in one of the comparisons and was supported by at least one of the other four studies.

### 2.12 Cytokine array assay

Cytokine profiling was performed using Proteome Profiler^TM^ Array Human (XL) Cytokine Array kit (ARY022B-R&D Systems), which detects 105 different cytokines in duplicate. We used cell culture supernatants collected from glioma patient-derived cells grown in 2D, GSC, and 3D-GMH conditions. Cytokine arrays were incubated overnight at 4 degrees with 500µl of cell culture supernatant. Following incubation with antibody detection cocktail, antibody conjugation, and recommended washes, membrane immunoblots were developed with the Chemiluminescent Substrate Reagent Kit provided using ChemiDoc MP Imaging System. Cytokine images were analyzed using ImageJ to determine both signal and background intensities. For background correction, median background intensities were subtracted from mean signal intensities. The mean pixel density of the cytokine/chemokine spots of 3D-GMH were normalized to 2D values to calculate the differential expression.

### 2.13 InnateDB interaction analysis

InnateDB(29) is a database containing the innate immunity interactome generated through manual curation of several public databases. All human gene interactions with at least one cytokine identified with increased secretion in 3D-GMH vs. 2D conditioned media were selected. Darmanis et al. single-cell dataset was used to determine expression levels of each gene in immune cells from the periphery and immune cells from the core of the tumor. Ivy GAP data was used to determine if the gene is preferentially expressed in any spatial locations. Average log2 CPM values were calculated for each spatial location and then z-scored. The location with maximum z-score (z-score >1) was assigned to that gene. PAN and PNZ_ccGSC were grouped as Perinecrotic, MVP and HBV_ccGSC were grouped as Perivascular (PV), LE and IT as Tumor periphery, and CT and CT_control were grouped as Tumor core. The network was plotted using cytoscape to show the interactions of the cytokines with increased secretion in 3D-GMH vs. 2D conditioned medium(30).

### 2.14 Macrophage infiltration assay

The conditioned medium of cells cultured in 2D (at 70% confluency) and cells cultured in 3D-GMH (after 15 days) were collected. Both cell types were also cultured as neurospheres in NBM in serum-free conditions in order to enrich them for glioma stem-like cells(31), and the conditioned media was collected when the neurosphere turned slightly necrotic (between 6-8 days).

To study macrophage infiltration, U937 cells were cultured in RPMI1640 media and induced by 5nM PMA (phorbol 12-myristate-13-acetate) (P1585, Sigma-Aldrich) for 48 hours as per protocol(32) to differentiate them into macrophages. To check the level of infiltration of macrophages, a standard transwell Boyden chamber invasion assay was performed as per the protocol(33), where U937 derived macrophages 4*10^5^ in the top chamber migrated towards condition media in the bottom chamber, which constitutes secretome of GBM cells cultured on respective culture conditions.

### 2.15 Isolation of Human monocytes

Human peripheral blood was collected using sodium citrate as anticoagulant (heparin and EDTA have also been used as anticoagulants) and layered onto a cushion of Ficoll-Hypaque (1.077 g/ml) and centrifuged at room temperature for 30 minutes at 400g. The mononuclear cells were collected from the interface and washed twice with phosphate buffer, pH 7.4. These isolated cells were seeded into a T25 flask and incubated for 2 hours, adherent monocytes were separated from non-adherent lymphocytes and used for immune-infiltration assay.

## 3. Results

We used seven high-grade glioma samples (grade III & IV) (Supplementary File - 1) collected from patients who received a craniotomy at MSMC along with glioblastoma cell line U251 for our study. Patient-derived glioma cells were cultured either as neurospheres in serum-free media with growth factors to enrich for glioma stem-like cells (GSC)(22) or cultured in 2D, 2D under hypoxia, and 3D-GMH with serum. Previously, we have characterized 3D-GMH (10% GelMA) for its tunable mechanical property, it was found that they are viscoelastic in nature and are stiff with 4.81 ± 0.73 kPa and the stiffness increases to 19.25 ± 6.85 kPa for 15% GelMA hydrogels(15). As the tumor microenvironment of glioblastoma is viscoelastic in nature and the stiffness of the tumor tissue ranges from 1-13 kPa(34), we employed 3D-GMH to understand the property of glioblastoma cells in this 3D model. The 3D-GMH was also found to be porous and the pore size decreased with increase in GelMA concentration in a linear fashion(15). It was also found that the 3D-GMH were cell responsive as the cancer cells cultured on 3D-GMH were able to attach, form spheroids and were able to invade into the hydrogel by upto 500µm(15) proving to be suitable for our study. Cells grown under all three culture conditions at early passages were further used for functional assays as well as transcriptome profiling (Fig-1a).

**Figure 1:**
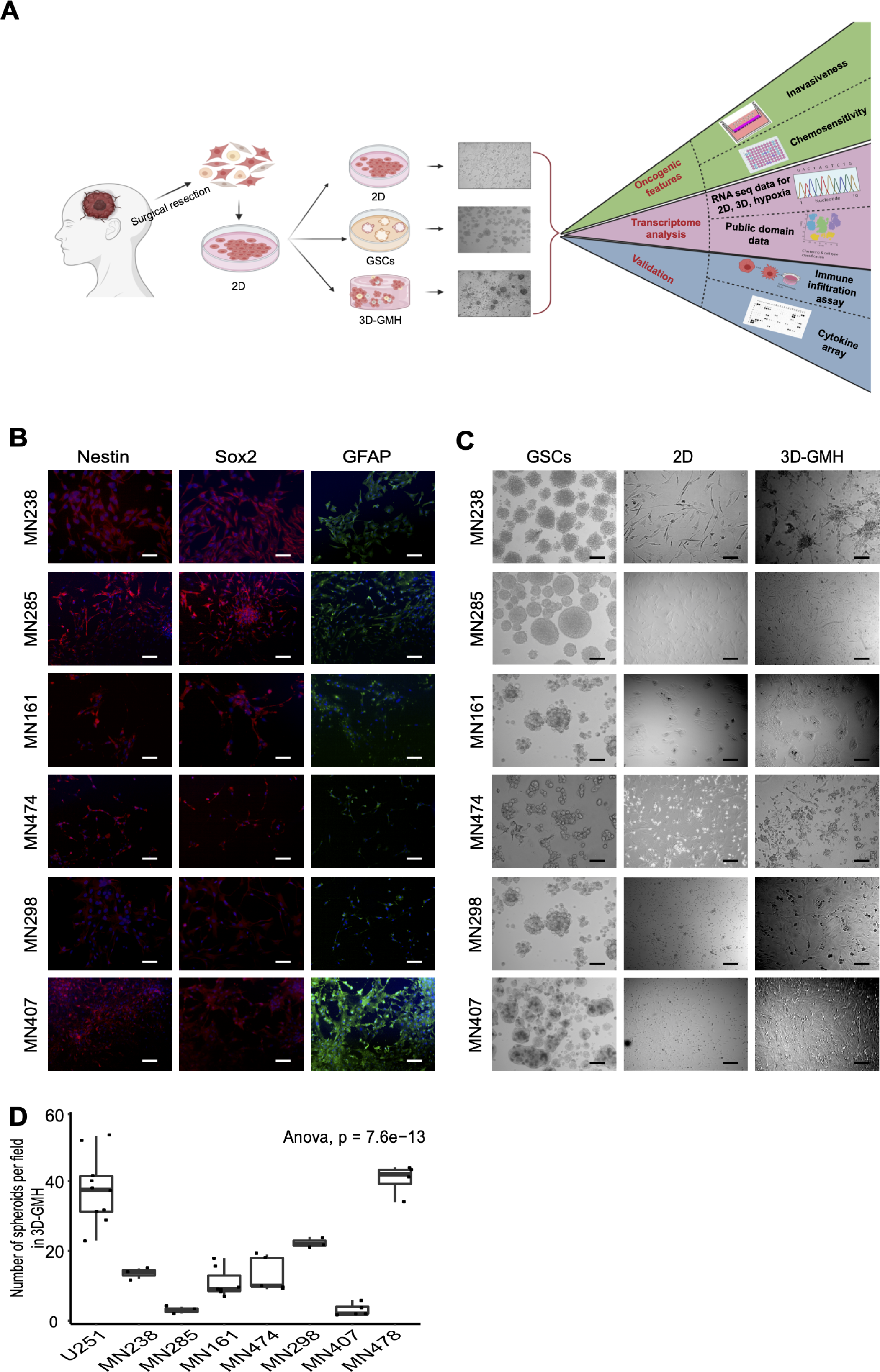
Study design, characterization, and phenotypic properties of glioma cells. (A) Schematic diagram of the experimental approach (created with BioRender.com) (B) Immunofluorescence analysis of patient-derived glioma cells for stem cell markers Nestin and SOX-2, and lineage marker GFAP. (C) Bright-field microscopy images depicting the morphology of patient-derived glioma cells grown in 2D, GSCs, and 3D-GMH. (D) Quantification of spheroid formation efficiency of patient-derived cells on 3D-GMH (ANOVA, p-val = 7.6e-13) Scale bar -100µm.

### 3.1 Primary glioma cells grown on GelMA hydrogels display heterogeneous morphology

Patient-derived glioma cells were characterized by immunofluorescence using Nestin, a neural progenitor marker, and SOX2, a marker for pluripotency and self-renewal. The lineage of the patient-derived glioma cells was confirmed by the expression of the astrocytic marker GFAP (Fig-1b). *In vitro* 3D cultures were established by culturing glioma cells in 3D-GMH for up to 15 days, with media changed every 3-5 days. Cells cultured in 3D-GMH adhered to the scaffolds within 24 hours. Within 5 days of culture these cells formed spheroids (Fig-1c) along with the characteristic prominent invadopodia-like structures to help them migrate out of the spheroids. Unlike the cells cultured in 3D-GMH, those cultured in 2D displayed a flattened spindle-like morphology (Fig-1c, Supplementary Figure - S1a)(35). Patient samples MN285 and MN407 gave rise to fewer spheroids, which may be attributed to patient-specific characteristics (Fig-1d).

### 3.2 Glioma cells cultured on GelMA hydrogels display variable phenotype with increased migratory potential, increased chemoresistance, and altered cell-cycle

To evaluate whether 3D-GMH could induce enhanced invasive properties in glioma cells, U251 and patient-derived glioma cells cultured in 3D-GMH were subjected to the standard trans-well Boyden chamber invasion assay using trans-well inserts coated with matrigel of 1mg/ml concentration. We observed that the U251 cells cultured in 3D-GMH displayed 3.5-fold higher invasive potential than their 2D counterparts. Similarly, MN238, MN298, and MN474 patient-derived glioma cells displayed 3, 3.5, and 7-fold higher invasive potential, respectively compared to the cells cultured in 2D (Fig-2a, 2b). U251 and MN238 cells cultured under GSC condition also displayed 2-fold higher invasive potential as compared to their 2D counterparts; however, it was lower compared to the 3D-GMH condition (Supplementary Figure - S2).

**Figure 2:**
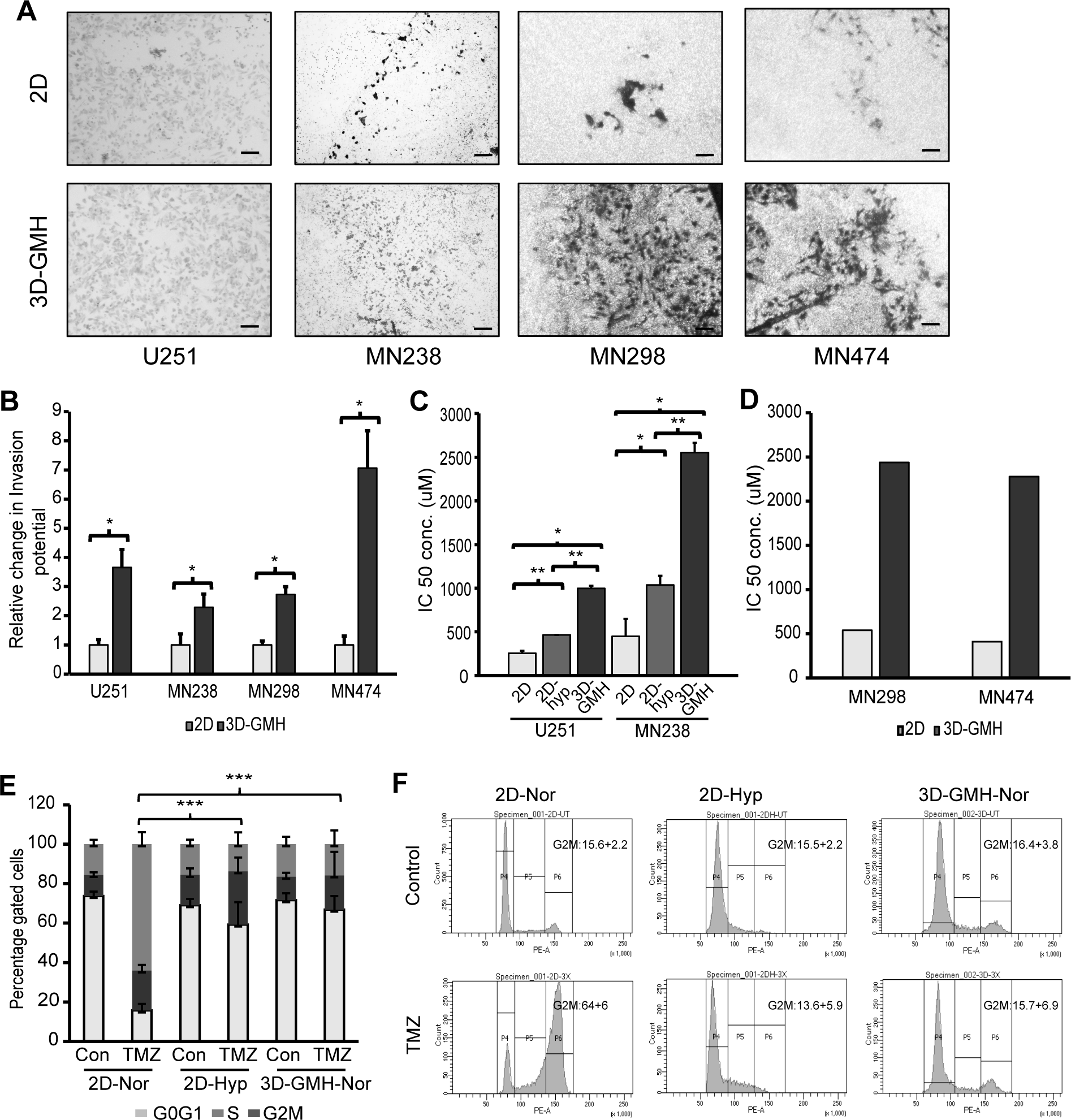
Culture on GelMA hydrogels enhances invasion and chemoresistance potential of glioma cells. (A) Boyden chamber invasion assay for U251 cells and patient-derived glioma cells cultured in 2D and 3D-GMH. (B) Quantification of the invaded cells. (C) Effect of TMZ on the viability of U251 and MN238 cells. U251 and MN238 cells incubated with various concentrations of TMZ were monitored using MTT assays after a 72-hour incubation period. (D) Effect of TMZ on the viability of MN298 and MN474 cells incubated with various concentrations of TMZ and monitored via MTT assays after a 72-hour incubation period. (E) Quantification of cell-cycle analysis of U251 cells cultured in 2D, 3D-GMH in normoxia, and in 2D hypoxia. Flow-cytometry analysis 72 hours after U251 cells were treated with a lethal concentration (3x of IC50) of TMZ revealed reduced G2/M arrest in cells cultured in 3D-GMH in normoxia and cells cultured in 2D hypoxia in comparison with cells cultured in 2D. (F) Representative flow cytometric DNA content of U251 cells labeled with Propidium Iodide in three independent experiments. The error bars represent the standard deviation. *indicates p<0.05, **indicates p<0.01, ***indicates p<0.001. (Abbreviations are Nor- normoxia (21% O2), Hyp-hypoxia (1% O2))

Further, to determine whether 3D-GMH can influence chemoresponse of glioma cells, we assessed their response to TMZ, an alkylating agent. U251 cells and patient-derived glioma cells were cultured in 3D-GMH for 5 days, followed by treatment with TMZ for 72 hours and MTT cell viability assay. In comparison to U251 and MN238 cells cultured on 2D (with IC50 of 254.3±29.43 µM and 448.1±201.5 µM respectely), we observed 4-6-fold increase in resistance to TMZ for both U251 and MN238 cells cultured in 3D-GMH with IC50 of 996.23±33.43 μM (p-val. < 0.001) and 2555±109.27 μM (p-val. < 0.001) (Fig-2c). Other patient-derived glioma cells MN298 and MN474 with IC50 of 538.1 μM and 410 μM respectively on 2D also showed a similar pattern with a 5-fold increase in the chemoresistance with IC50 of 2439 μM and 2278 μM respectively when cultured on 3D-GMH (Fig-2d). Hypoxia has previously been shown to influence chemoresponse to TMZ(36). Therefore, we wanted to determine whether the increased resistance to TMZ could simply be explained by the hypoxic conditions created when these cells migrate into the scaffold. So, to assess the role of hypoxia, both U251 and MN238 cells were cultured in 2D under hypoxic conditions (1% O2). U251 cells and MN238 patient-derived cells cultured in 2Dhypoxia showed IC50 values that were 2-fold higher with IC50 of 464.06±21.71 μM and 1036.77±105.14 μM respectively, as compared to cells cultured in 2D under normoxia (21% O2) (p<0.05). This demonstrates that hypoxic conditions does indeed affect cells’ resistance to TMZ. However, the IC50 value of these when cultured in 3D-GMH was 2-3-fold higher as compared to the IC50 of the cells cultured under 2D hypoxia (Fig-2c). Further, we performed cell cycle analysis to evaluate the drug-induced cytotoxicity in these cells. We observed that U251 cells cultured in 2D in normoxia resulted in an efficient G2-M phase arrest upon treatment with the lethal concentration of TMZ (3 times the concentration of IC50). U251 cells cultured in both 2D hypoxia and 3D-GMH did not display any G2-M phase arrest upon treatment with the lethal concentration of TMZ (Fig-2e, 2f), confirming that both 3D-GMH culture and hypoxia influences cells’ response to chemotherapeutic drugs. Taken together, we concluded that the glioma cells cultured in 3D-GMH acquire an increased TMZ-resistant phenotype, which could not entirely be explained by hypoxia.

### 3.3 Glioma cells grown on GelMA hydrogels resemble neoplastic cells found around hypoxic and vascular regions of the tumor

We generated RNA-seq data for patient-derived glioma cells grown under three different conditions; (1) 2D – grown in 2D with 10% serum, (2) 3D-GMH – grown in 3D-GMH with 10% serum, and (3) 2D hypoxia – grown in 2D with 10% serum in the hypoxia compared to normoxia. We used publicly available RNA-seq data for cells grown as neurospheres in serum-free conditions (GSCs) and differentiated glioblastoma cells (DGCs) in 2D with serum conditions(10,26–28). We performed a total of 18 comparisons (three 3D-GMH vs. 2D, one 3D-GMH vs. 2D – passage >5, two 3D-GMH vs. 2D hypoxia, two 3D-GMH vs. 2D hypoxia – passage >5, two 2D hypoxia vs. 2D, one 2D hypoxia vs. 2D – passage >5, seven GSCs vs. DGCs) (Supplementary File - 2) (Supplementary Figure – S3). We compiled two gene lists, one for genes overexpressed under 3D- GMH culture conditions compared with 2D and another for genes overexpressed under neurosphere conditions (GSC) compared to the cells that were allowed to differentiate under 2D serum conditions (DGC).

Neftel and co-workers generated and analyzed single-cell RNA-seq data from 28 pediatric and adult glioblastoma tumors to identify four major neoplastic cell-types defined by six gene modules; (1) Mesenchymal – Hypoxia independent (MES1like) and hypoxia dependent (MES2like) mesenchymal related gene sets, (2) Astrocytic - astrocytic (AC-like) marker gene set, (3) Oligo- oligodendroglial (OPC-like) lineage marker gene set, and (4) Neural-stem and progenitor cell signatures (NPC1-like and NPC2-like), as well as two cell cycling modules (G1S and G2M - CC) (Supplementary File - 3). We wanted to understand the anatomic locations where each of these cell-types reside. The Ivy Glioblastoma Atlas Project (Ivy GAP) generated molecular signatures of cells present in five major anatomic features of glioblastoma, visible by H&E staining, i.e., leading edge (LE), infiltrating tumor (IT), microvascular proliferation (MVP), cellular tumor (CT), and pseudopalisading cells around necrosis (PAN). In addition to this, the atlas also contains transcriptome profiles of cells expressing community-curated glioblastoma stem cell associated genes (ccGSC markers) (Table 1). These samples were grouped into three categories; (1) CT area with expression of one or more ccGSC markers (CT_ccGSC), (2) Perinecrotic zone with expression of one or more ccGSC markers (PNZ_ccGSC), and (3) Area around hyperplastic blood vessels with expression of one or more ccGSC markers (HBV_ccGSC). The CT_control samples are CT regions with the absence of one or more ccGSC markers. Single sample gene set enrichment analysis (SSGSEA) using Neftel et al. gene modules shows that MES1-like signature is enriched in PAN, PNZ_ccGSC, CT_ccGSC, and HBV_ccGSC regions whereas MES2-like signature is enriched in mostly PAN and PNZ_ccGSC regions (Fig-3a). CC signature is enriched in all regions except LE, PAN, and PNZ_ccGSC. The AC-like and OPC-like cell-type signatures are enriched in LE, IT, CT, and CT_control. NPC1-like cell-type signature is enriched in IT, CT, and CT_control whereas NPC2-like signature is restricted to LE, and IT regions. We wanted to understand the cell-type and the anatomic feature best mimicked by cells grown in 3D-GMH and GSC conditions by analyzing the shift in their transcriptional profiles. We took all the genes that were differentially expressed in more than one comparison. The data showed that genes overexpressed in each of the 3D-GMH vs. 2D comparisons are enriched in MVP, PAN, CT_ccGSC, HBV_ ccGSC, and PNZ_ ccGSC. In contrast, genes overexpressed in GSC vs. DGC comparisons are overexpressed in IT, CT, and CT_control. Of note, the overexpressed genes in cells grown under hypoxia show inconsistent enrichment for IT, PAN, and PNZ_ccGSC (Fig-3b). This may be due to the differences among the patient-derived cells that impact their response to hypoxic conditions. The compiled 3D-GMH gene list showed enrichment in MVP, HBV_ccGSC, and PNZ_ccGSC, whereas the GSC gene list shows enrichment in the IT, CT, and CT_control regions (Fig-3c). We also compared the compiled 3D-GMH and GSC gene lists with the clusters of single-cell glioblastoma data(8, 37). Each single cell cluster of neoplastic cells was assigned the cell subtype as described earlier(8, 25). Genes in the 3D-GMH list are overexpressed in mesenchymal cell-types found in hypoxic and vascular regions, whereas genes in the GSC list are overexpressed in AC-like, OPC-like, and CC cell-types found in IT and CT regions (Fig-4a). We also performed GO enrichment analysis and found biological processes related to immune response, cytokine production, and cell motility enriched in 3D-GMH in contrast to nervous system development related processes enriched in the GSC list (Fig-4b). Given the cells grown in 3D-GMH resembled the stem-like mesenchymal cells around perinecrotic and perivascular zones and are known to be more resistant to treatment(38), we hypothesized that the overexpression of genes in the 3D-GMH list would be associated with poor survival. Using TCGA and CGGA data, we found that the patients with lower expression of genes in the 3D-GMH list and higher expression of genes in the GSC list had significantly better survival than patients with higher expression of genes in 3D-GMH list and lower expression of genes in GSC list (Fig-4c, Supplementary File - 4). Overall, these results demonstrated that the glioma cells grown in 3D- GMH enrich for mesenchymal-like cells, possibly due to the mechano-biological cues from the 3D-GMH microenvironment.

**Figure 3:**
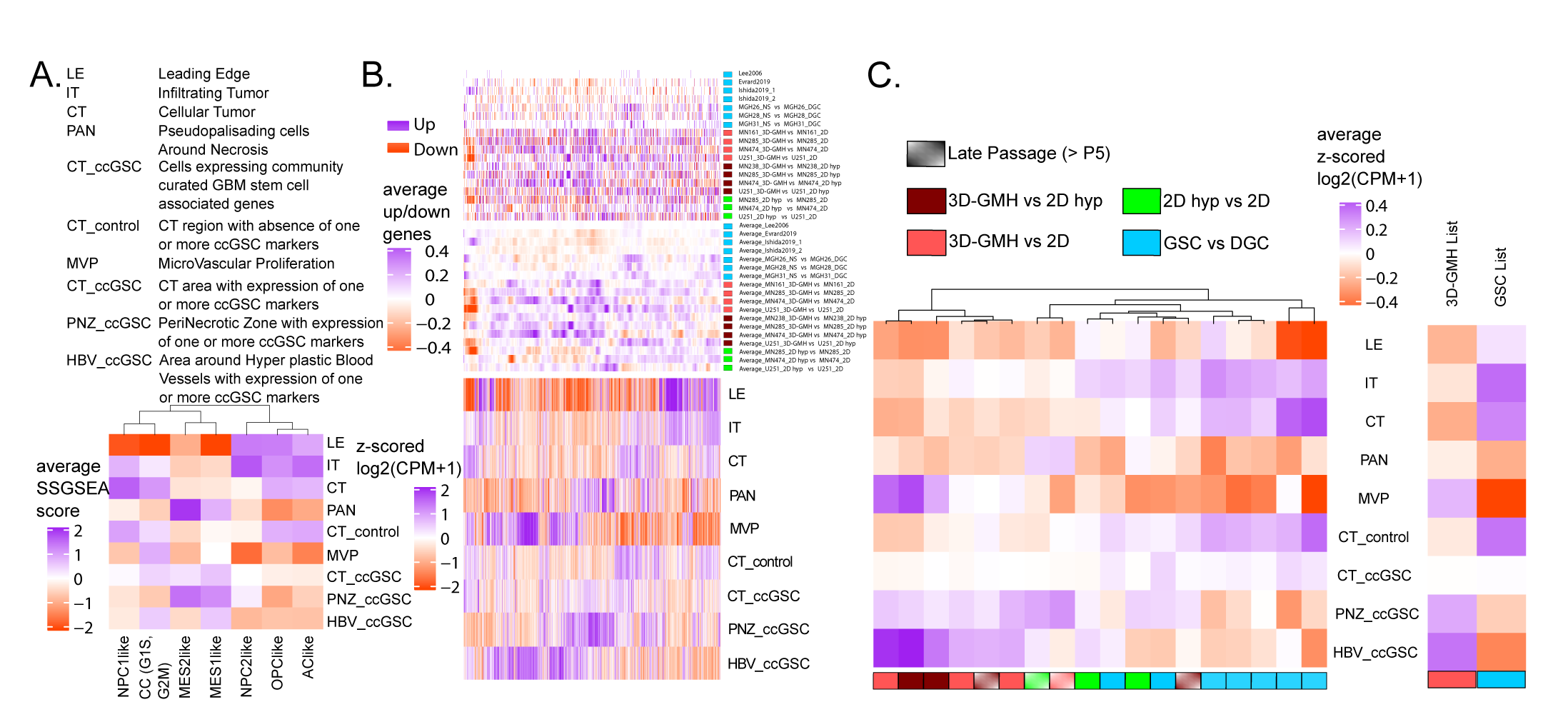
Transcriptional profiles of cells grown on GelMA hydrogels resemble the profiles of cells in perinecrotic and perivascular zones of tumor. (A) The columns represent the glioblastoma cell-types (meta-modules as described in Neftel et al.) and the rows represent the Ivy GAP RNA-seq data consisting of spatial transcriptomes based on histology and expression of community curated glioblastoma stem cell-associated genes (Table 1). Single sample geneset enrichment analysis (SSGSEA) scores(52) for each meta-module were averaged over each Ivy GAP spatial feature. Mesenchymal cell-type signatures (MES1-like,MES2-like) are enriched in PAN, PNZ_ccGSC, CT_ccGSC, and HBV_ccGSC regions. Cell cycling cell-type signature (G1S, G2M) is enriched in all regions except LE, PAN, and PNZ_ccGSC. The astrocyte and oligo-like cell-type signatures (AC-like, OPC-like) are enriched in LE, IT, CT, and CT_control. Neural progenitor-like cell-type signatures (NPC1, NPC2) are enriched in LE, IT, CT, and CT_control. (B) Ivy GAP expression matrix for differentially expressed genes in at least two out of 18 comparisons (Supplementary Figure - S3, Supplementary methods). The genes are hierarchically clustered. The top panel shows which gene is up (purple) and down (orange) regulated for each condition. The second panel shows the average of the top panel over 100 genes. We can see that genes overexpressed in each of the 3D-GMH vs. 2D comparisons are enriched in MVP, PAN, CT_ccGSC, HBV_ ccGSC, and PNZ_ ccGSC. In contrast, genes overexpressed in GSC vs. DGC comparisons are overexpressed in IT, CT, and CT_ control. (C) Ivy GAP expression matrix averaged over each spatial feature for a set of upregulated genes. Columns represent each of the 18 comparisons and are clustered. 3D-GMH vs. 2D comparisons cluster together with enrichment in PNZ_ccGSC and HBV_ccGSC spatial features in contrast with GSC vs. DGC comparisons that show enrichment in IT, CT, and CT_control features. The combined list of consistently overexpressed genes in 3D-GMH vs. 2D comparison (3D-GMH List) and GSC vs. DGC comparisons (GSC List) show a similar trend.

**Figure 4:**
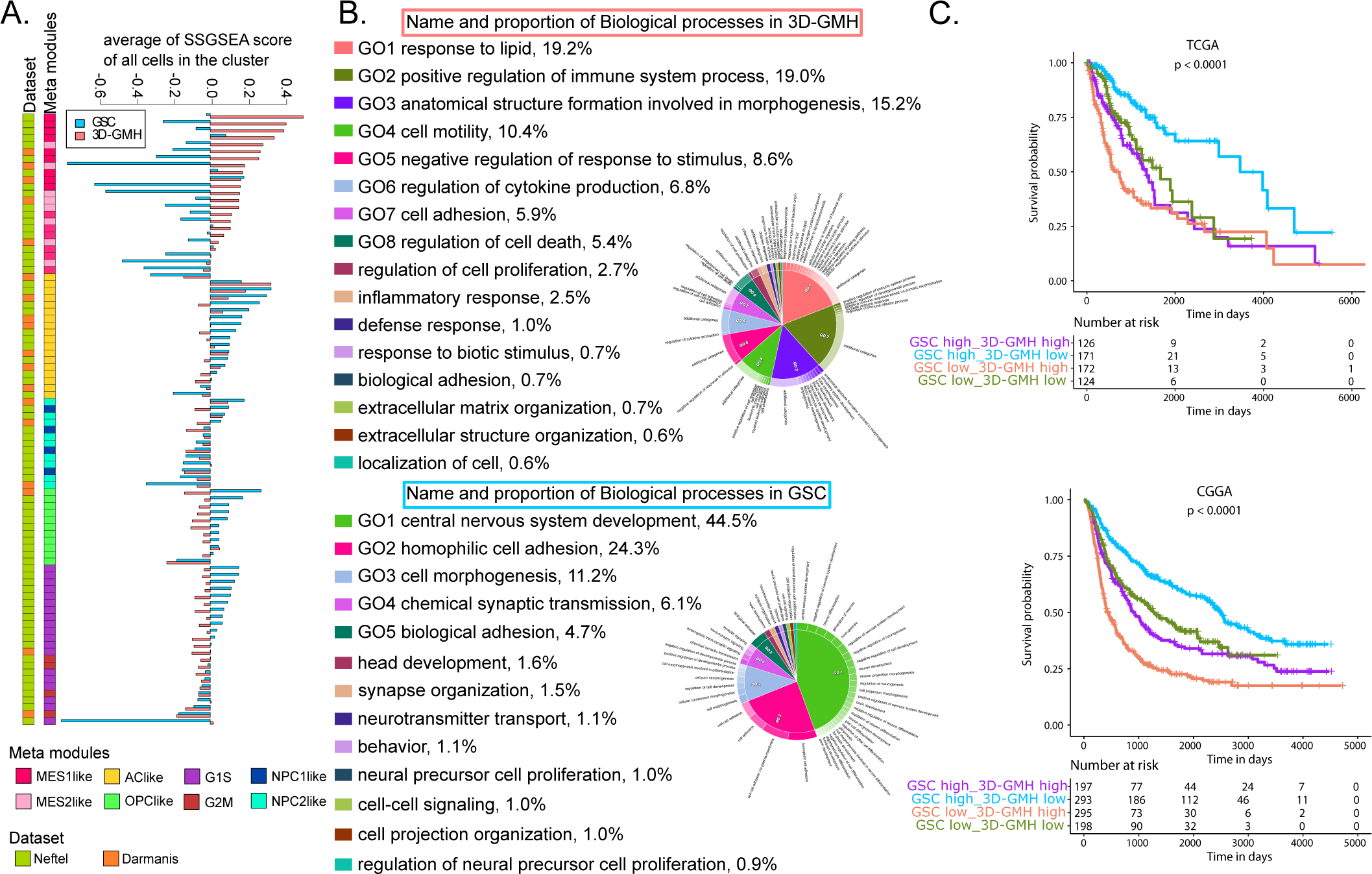
Glioma cells cultured on GelMA hydrogels resemble mesenchymal-like glioblastoma cells associated with immune response and poor survival. (A) 3D-GMH and GSC gene list SSGSEA score comparisons across clusters from the Darmanis et al. and Neftel et al. single-cell datasets(44). Only the neoplastic cell clusters were selected for this analysis. Labels were assigned to all clusters based on the cell-type which has maximum frequency in that cluster. The genes in the 3D-GMH list are overexpressed in mesenchymal cell- types, whereas genes in the GSC list are overexpressed in AC-like, OPC-like, and cell cycling cell- types. (B) Gene ontology enrichment analysis using Toppfun(53) for the 3D-GMH and GSC gene lists. The plots were made using CirGo tools(54). The 3D-GMH list showed enrichment of biological processes like response to lipid, cell motility, and regulation of immune system processes, whereas the GSC list showed enrichment of processes like central nervous system development and cell adhesion. (C) Survival analysis of TCGA [https://portal.gdc.cancer.gov/] and CGGA[http://www.cgga.org.cn/] datasets using 3D-GMH and GSC lists. Based on the median SSGSEA score of 3D-GMH and GSC gene lists, we divided the glioma samples into four categories in TCGA and CGGA datasets. The patients with lower SSGSEA score for 3D-GMH list and higher score for GSC list have significantly better survival compared to patients with higher score for 3D-G:MH list and lower score for GSC list (median survival of 3460 days, 95% CI [2988, NA] vs. median survival of632 days 95% CI [537, 1062] in TCGA and median survival of2499 days 95% CI [2237, 2982] vs. median survival of 432 days 95% CI [379, 567] in CGGA).

### 3.4 GelMA hydrogels enhance the ability of glioblastoma cells to recruit macrophages

Tumor-associated mesenchymal stem cells are known to facilitate tumor growth by releasing growth factors and cytokines(39). Considering the enrichment of mesenchymal-like cells in 3D- GMH, we sought to characterize the 3D-GMH secretome. Using cytokine arrays, conditioned media (CM) of glioma cells grown in 2D, GSC, and 3D-GMH were analyzed. (Fig-5a, 5b, Supplementary Figure - S4). In agreement with our RNA-seq analysis, 3D-GMH secretome analysis showed increased secretion of tumor growth/invasion markers, pro-inflammatory cytokines, pro-angiogenic factors, and complementary factors (Fig-5a, 5b), suggesting that 3D- GMH condition enhanced the secretion of pro-tumorigenic cytokines.

**Figure 5:**
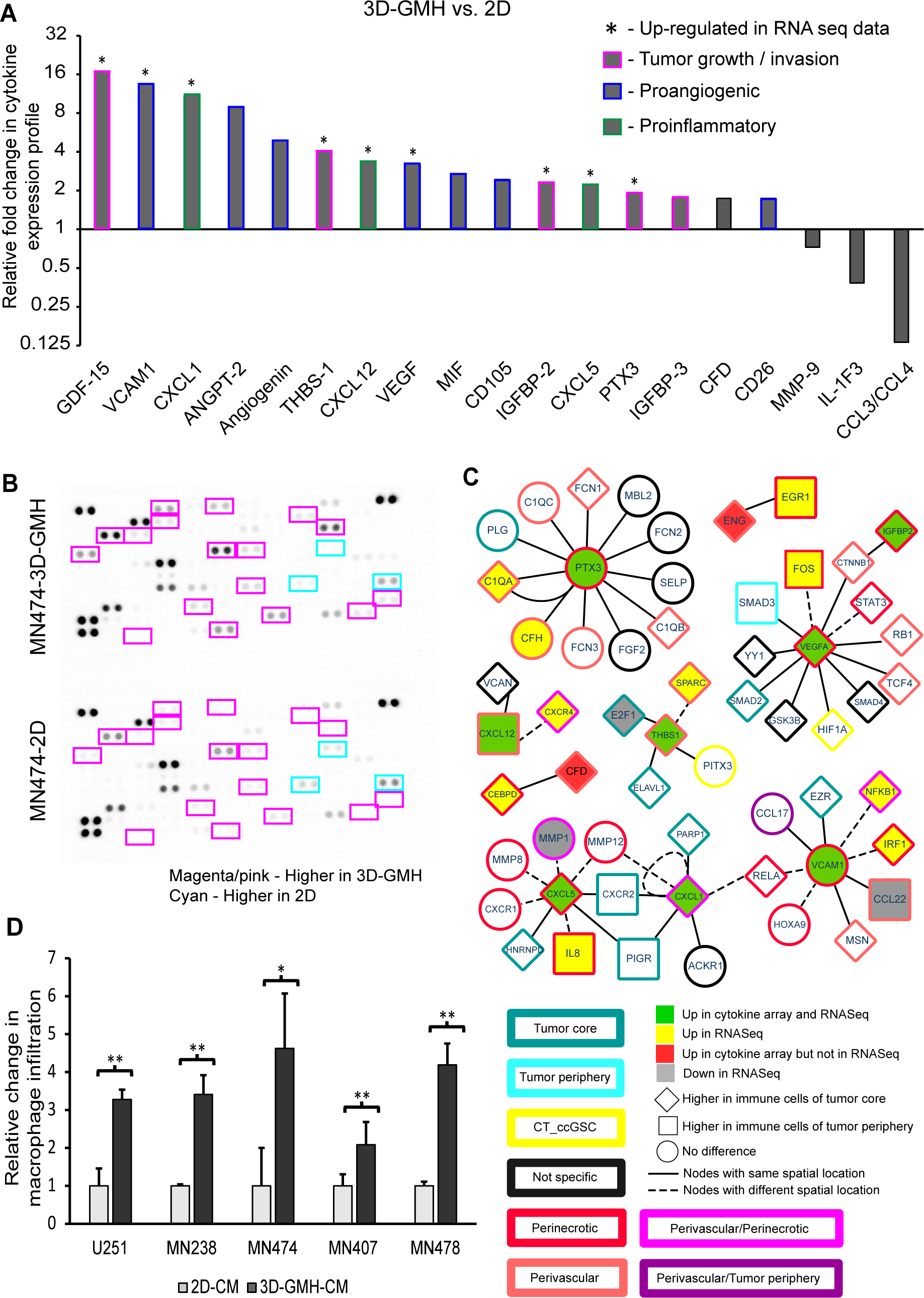
Glioma cells cultured on GelMA hydrogels enhance immune cell recruitment. (A) Fold change in cytokine RNA expression under 3D-GMH relative to 2D. Mean pixel density is normalized to 2D, + indicates that the respective gene is upregulated in RNA-seq data. (B) Cytokine levels in the CM of MN474 glioma cells cultured in 3D-GMH (Top) or 2D (Bottom). Magenta/pink bordered cytokine indicates enhanced secretion in cells cultured in 3D-GMH, and Cyan bordered cytokine indicates enhanced secretion in cells cultured on 2D. (C) The network derived from InnateDB(29) for cytokines that are enhanced in 3D-GMH vs. 2D CM generated using Cytoscape(30). The fill color indicates the differential expression of the gene in the cytokine array as well as RNA-seq data (MN474 3D-GMH vs. MN474 2D). The shape indicates if the gene is differentially expressed in immune cells of the tumor core vs. tumor periphery(37). The border color shows the spatial location where the gene is preferentially expressed. The error bar represents the standard deviation. *indicates p<0.05, **indicates p<0.01. (D) Boyden chamber invasion assay for U937-derived macrophages against CM of U251 cells and patient-derived glioma cells cultured in 2D and 3D-GMH.

Further, the functional significance of possible molecular interactions of upregulated cytokines in the RNA-seq and cytokine arrays was assessed using InnateDB(29). Network analysis revealed that most of the genes enriched in 3D-GMH interact with the secreted cytokines, expressed in the perivascular/perinecrotic zone of the tumor core and in the immune cells of the tumor core (Fig-5c). Molecular links showed that interactions of VEGFA with IGFBP2 activate other angiogenesis factors facilitating angiogenesis and neovascularization(40). Also, inflammatory factors like CXCL1, CXCL5, CXCL12, CXCR4, VCAM1, and IL8 interact with matrix metalloproteinases(41, 42), and association of THBS1 with PITX3, SPARC(43) plays a significant role in enhancing the recruitment of macrophages. Furthermore, hypoxia-regulated proteins interact with ficolins and complement factors to activate the complement pathway(44, 45), thereby playing an essential role in the inflammatory process and attracting immune cells. Together, 3D- GMH secretome showed an increase in cytokines involved in tumor growth, angiogenesis, invasion, pro-inflammatory cytokines, and complement factors majorly involved in recruiting immune cells.

Thus, to determine whether glioma cells cultured in 3D-GMH can recapitulate the immune cell- infiltration behavior, we used a PMA-induced U937- macrophage cells in a standard Boyden chamber invasion assay. The U937 derived macrophages were seeded in the Boyden chamber’s top compartment with the CM of 2D, 3D, and neurospheres (NS) in the bottom chamber and compared invasion of macrophage cells to the CM of 2D, GSC, and 3D-GMH. Despite variations between samples, which can be attributed to patient characteristics, we observed that macrophage infiltration was highest under the influence of 3D-GMH-CM (minimum of 3-fold), GSC-CM (2-fold) compared to 2D-CM (Fig-5d, Supplementary Figure - S5a). A similar pattern was also observed with human monocytes isolated from a healthy individual (Supplementary Figure - S5b, S5c), suggesting that glioma cells cultured in 3D-GMH have enhanced ability to recruit immune cells.

In summary, our study shows that glioma cells grown in 3D-GMH exhibit mesenchymal-like characteristics with enhanced cytokine production and partially recapitulate the immune cell- infiltration.

## 4. Discussion

In spite of a plethora of treatment approaches with promising results in preclinical studies being tested in human clinical trials, only one new treatment has been recently approved for glioblastoma(46), thus highlighting the need for new *in vitro* models as a bridge to *in vivo* studies. Given that cellular and microenvironmental heterogeneity is considered the main cause of treatment failure(1,2,5), we hypothesized that the traditional 2D serum-based monolayer and serum-free GSC generating culture conditions are insufficient to capture quiescent mesenchymal glioma cell-types(5). It is important to develop cell culture model for growing these mesenchymal cell-types that have escaped *in vitro* studies so far. This will allow for engineering of biomimetic models suitable for drug screening with a higher hit rate. Glioma cells cultured in 3D systems that provide more realistic cell-cell and cell-matrix interactions exhibit changes in cellular morphology, phenotype, gene expression pattern, and drug resistance(47, 48). In this study, through integrated transcriptomic analysis of bulk, single-cell, and spatial RNA-seq data, we show that patient- derived glioma cells grown under GSC conditions resemble oligo-, neural-, and astro-like glioma cells, whereas cells grown in 3D-GMH resemble the mesenchymal-like glioma cells.

Single-cell transcriptome analysis of isocitrate dehydrogenase wildtype (IDHwt) glioblastoma and IDH-mutant astrocytomas revealed different patient-independent glioma cell-types, i.e., OPC-like, NPC-like, Mes-like, AC-like, and cell cycling stem cell-like(8, 49). Wang and co-workers showed that the GSCs from IDHwt glioblastoma cells lie on a single gene signature axis ranging from being proneural (highly proliferative) to mesenchymal phenotype (quiescent and a secretory phenotype)(9). Overlaying single-cell transcriptome data on the anatomic feature transcriptome atlas(1) shows that mesenchymal-like cells reside in hypoxic and perivascular niches, whereas the other cell-types are found more in the infiltrative edge. Krieger and co-workers used a co-culture system where they grew glioblastoma cells (GSCs) with human induced pluripotent stem cell- derived brain organoids. Cells grown in monoculture showed an astrocytic signature, which shifted to NPC-like signature when co-cultured, indicating that the model mimics the infiltrative edge(50). Another study used hyaluronic acid-rich hydrogels of different stiffness and four cell-types (GSCs, macrophages, astrocytes, and neural stem cells) for a 3D bioprinted model to mimic glioblastoma- immune cell interactions(18). This model too showed increased expression of genes OLIG2 and ki67 which are markers of proneural GSCs and absent in mesenchymal-like cells. Pine and co- workers extensively compared and contrasted patient-derived cells grown in 2D, matrigel, cerebral organoids, and xenograft tumors using single-cell sequencing (45). These culture systems did not show enrichment of hypoxia-independent mesenchymal cells. One of the possible reasons is use of serum-free neurobasal medium in all the recent studies of 3D cell culture systems for glioblastoma. Our study shows the cells grown in 3D-GMH with serum overexpress and secrete CXCL1, CXCL5, IGFBP2, PTX3, THBS1, VEGFA, and VCAM1 - all genes expressed in perivascular or perinecrotic zone of glioblastoma tissues that are hotspots for immune cells(5). Many of these factors are known to enhance the recruitment of immune cells(41–44), and we see this in our assays where 3D-GMH CM attracts a higher number of immune cells compared to 2D and GSC CM. Now, from single-cell data, we know that the immune cells found in the tumor periphery are different from the ones present in the core of the tumor(37). When we analyzed the interactors of these proteins that are enhanced in the 3D-GMH secretome(29), we found these were overexpressed on the immune cells found in the core compared to the periphery of the tumor.

## 5. Conclusion

Culturing glioma cells in 3D-GMH captures cellular phenotypes found in perivascular and perinecrotic zones. Our study shows for the first time that the use of a 3D hydrogel along with serum to grow early passage patient-derived cells allows us to culture hypoxia-independent glioma cells. In the future, we expect that our approach will include glioma samples from a much wider spectrum. Each anatomic feature of glioblastoma consists of specific composition of cell-types like immune cells, vascular cells, astrocytes, oligodendrocytes, neurons, etc., in addition to the neoplastic cells. For example, perivascular niche contains endothelial cells, immune cells, and hypoxia-independent mesenchymal-like cells, whereas perinecrotic zone consists of immune cells and hypoxia-independent and dependent mesenchymal-like cells(51). Enriching the complexity of the hydrogel by incorporating various extracellular matrix (ECM) molecules specific to brain tissue like hyaluronic acid, by varying the mechanical properties of the hydrogel, and by co-culturing multiple cell-types will help us in modeling specific niches; perivascular vs. perinecrotic(18).

## Disclosure

The authors declare no conflict of interest

## Data availability

Processed expression count data is available through (https://github.com/MSCTR/3Dgelma). Further information and request for resources, raw data, and reagents should be directed to NS and AC.

## Author contributions

NS and AC designed the study. PMH and RAG performed the experiments. PMH prepared GelMA hydrogels and performed invasion, chemoresponse and immune infiltration assays. RAG and DN prepared patient samples, RAG performed IFC and cytokine array experiments. NS and PS performed bioinformatic analysis. KPC performed surgeries and collected patient samples. AC and NS designed the experiments, helped with data analysis and interpretation of the data. All authors contributed towards writing and have read and approved the final version of the manuscript.

## Acknowledgments

We thank Dr. Paturu Kondaiah, IISc, for providing U251 cell line, Yogesh Pasupathy and Anurag CN for sample collection.

## Supplementary Information

Supplementary Figure legends

Supplementary Figure-S1: Phenotypic properties of glioma cells cultured in 2D and in 3D-GMH.

Supplementary Figure-S2: Glioma cells cultured as neurospheres show increased invasive property. Supplementary Figure-S3: Volcano plots representing differentially expressed genes.

Supplementary Figure-S4A-D: Glioma cells cultured in 3D-GMH secrete enhanced cytokine and chemokines. Supplementary Figure-S5A-C: Glioma cells cultured in 3D-GMH enhances primary monocyte infiltration.

Supplementary Files

Supplementary File -1: Representative patient-derived glioma and glioblastoma cell lines cultured under different conditions. Supplementary File -2: Gene matrix indicating the differential genes across 14 comparisons. Supplementary File -3: Neftel meta-modules genesets used for SSGSEA. Supplementary File -4: Survival data.

